# Extracellular Matrix Dysfunction in Sorsby Patient-Derived Retinal Pigment Epithelium

**DOI:** 10.1101/2021.01.06.425613

**Authors:** Abbi L. Engel, YeKai Wang, Thomas H. Khuu, Emily Worrall, Megan A. Manson, Rayne R. Lim, Kaitlen Knight, Aya Yanagida, Jian Hua Qi, Aravind Ramakrishnan, Richard G Weleber, Michael L. Klein, David J. Wilson, Bela Anand-Apte, James B. Hurley, Jianhai Du, Jennifer R. Chao

## Abstract

Sorsby Fundus Dystrophy (SFD) is a rare form of macular degeneration that is clinically similar to age-related macular degeneration (AMD), and a histologic hallmark of SFD is a thick layer of extracellular deposits beneath the retinal pigment epithelium (RPE). Previous studies of SFD patient-induced pluripotent stem cell (iPSC) derived RPE differ as to whether these cultures recapitulate this key clinical feature by forming increased drusenoid deposits. The primary purpose of this study is to examine whether SFD patient-derived iPSC-RPE form basal deposits similar to what is found in affected family member SFD globes and to determine whether SFD iPSC RPE may be more oxidatively stressed. We performed a careful comparison of iPSC RPE from three control individuals, multiple iPSC clones from two SFD patients’ iPSC RPE, and post-mortem eyes of affected SFD family members. We also examined the effect of CRISPR-Cas9 gene correction of the S204C *TIMP3* mutation on RPE phenotype. Finally, targeted metabolomics analysis with liquid chromatography and mass spectrometry analysis and stable isotope-labeled metabolite analysis was performed to determine whether SFD RPE are more oxidatively stressed. We found that SFD iPSC-RPE formed significantly more sub-RPE deposits (∼6-90 μm in height) compared to control RPE at 8 weeks. These deposits were similar in composition to the basal laminar drusen found in SFD family member globes by immunofluorescence staining and TEM imaging. S204C *TIMP3* correction by CRISPR-Cas9 gene editing in SFD iPSC RPE cells resulted in significantly reduced basal laminar and sub-RPE calcium deposits. We detected a ∼18-fold increase in TIMP3 accumulation in the extracellular matrix (ECM) of SFD RPE, and targeted metabolomics showed that intracellular 4-hydroxyproline, a major breakdown product of collagen, is significantly elevated in SFD RPE, suggesting increased ECM turnover. Finally, SFD RPE cells have decreased intracellular reduced glutathione and were found to be more vulnerable to oxidative stress. Our findings suggest that elements of SFD pathology can be demonstrated in culture which may lead to insights into disease mechanisms.

## 1. Introduction

Sorsby Fundus Dystrophy (SFD) is a rare form of autosomal dominant macular degeneration that shares similar clinical features with age-related macular degeneration (AMD). Both SFD and AMD patients develop macular drusen, geographic atrophy, choroidal neovascularization, and progressive central vision loss (Anand-Apte et al., 2019; Weber et al., 1994). Individuals with SFD often experience an earlier onset of symptoms compared to their AMD counterparts, which results in central vision loss by as early as the fourth decade of life. While the pathogenesis of AMD is multifactorial, SFD results from mutations in a single gene, called tissue inhibitor of metalloproteinase 3 (*TIMP3*).

TIMP3 is secreted by the retinal pigment epithelium (RPE), and it plays a critical role in extracellular matrix (ECM) remodeling. It inhibits several matrix metalloproteinases (MMP-1, - 2, -3, and -9), a disintegrin and metalloproteinases (ADAMs), and ADAMs with thrombospondin motifs (Arpino et al., 2015; Stohr and Anand-Apte, 2012). TIMP3 also inhibits angiogenesis and inflammation by influencing VEGF-VEGFR2 interactions (Qi and Anand-Apte, 2015). While all TIMP family members are secreted, only TIMP3 is sequestered in Bruch’s membrane (BrM), a thin layer interposed between the RPE and underlying choriocapillaris (Jackson et al., 2017).

Bruch’s membrane is a semi-permeable barrier that provides structural support for the RPE and allows for nutrient flow from the choriocapillaris. A histological hallmark of SFD is the presence of a grossly thickened and disorganized BrM in post-mortem globes (Kuntz et al., 1996). The thickened BrM is hypothesized to function as a diffusion barrier preventing nutrients from reaching the RPE and retina from the choriocapillaris. SFD patients given high dose vitamin A supplements to counteract this were noted to have some restoration of night vision, albeit at an unsustainably high dose (Jacobson et al., 1995).

While mutant TIMP3 expression is suspected to play a role in the progressive thickening of BrM in SFD and subsequent RPE degeneration, the sole murine model, S179C *TIMP3*, does not reproduce these salient characteristics (Weber et al., 2002). This has made SFD an attractive candidate disease for study using patient induced pluripotent stem cell (iPSC)-derived RPE due to its early-onset and clinically severe phenotype attributable to a single gene (Galloway et al., 2017; Hongisto et al., 2020). However, findings differ as to whether SFD iPSC RPE recapitulate important features of the disease in culture, including increased drusenoid deposits or disrupted extracellular matrices. Recently, it was shown that the S179C *TIMP3* mouse model may be more susceptible to retinal degeneration after exposure to oxidative stress (Wolk et al., 2020). However, it has not been similarly shown whether human SFD RPE may have increased susceptibility to oxidative stress or alterations in metabolism. The availability of familial SFD post-mortem globes and generation of CRIPSR-corrected isogenic SFD iPSC lines in this study allows for comparative characterization of SFD iPSC-derived RPE phenotype in culture. Separately, because ECM dysfunction can impact cellular metabolism and susceptibility to oxidative damage (Bonnans et al., 2014), metabolic changes in patient-derived SFD RPE and effects on RPE survival under conditions of oxidative stress in culture are also examined.

## 2. Materials and Methods

### 2.1. Histology of post-mortem globes

Sorsby fundus dystrophy (SFD) globes were generously donated to the Lions Eye Bank of Oregon. Normal adult human globes were obtained from the University of Washington Tissue Bank for Ophthalmology Research (HSD# 41470). Standard protocols for hematoxylin and eosin (H&E), periodic acid-Schiff (PAS) and Movat’s pentachrome staining were performed on 5 μm-thick tissue sections. Images were taken at 40X on a Nikon Eclipse E1000 wide field microscope.

### 2.2. Clinical imaging

Informed consent was obtained from all subjects prior to inclusion in the study (University of Washington IRB-approved STUDY00010851), and experiments were conducted according to the principles expressed in the Declaration of Helsinki. Study participants underwent 30° digital fundus color imaging (Zeiss Visucam, Zeiss, Oberkochen, Germany) and optical coherence tomography imaging (Spectralis HRA-OCT; Heidelberg Engineering, Heidelberg, Germany).

### 2.3 Generation of patient induced pluripotent stem cells (iPSC)

iPSCs were generated as described previously (Du et al., 2016). In brief, PBMCs separated using a Ficoll-Hypaque gradient were cultured and reprogrammed via transfection of episomal vectors (OCT4, SOX2, KLF4, LIN28A, LMYC) and a short hairpin RNA targeting p53 using the Nucleofector II device. Transfected erythroblasts were plated onto irradiated mouse embryonic fibroblasts (iMEFs) in reprogramming medium until iPSC colonies emerge, then switched to hESC medium. iPSC colonies were picked and expanded first on iMEF plates, then on feeder-free plates. Cells were cultured in either E8 medium (Life Technologies, Carlsbad, CA, USA) or mTeSR (StemCell Technologies, Vancouver, BC, Canada). Karyotyping was performed and iPSC lines of normal karyotype were selected for study (Supplementary Fig. 1), and all iPSC lines were screened for AMD risk SNPs (Supplementary Table 1).

### 2.4. RPE differentiation and culture

iPSCs forming medium size colonies were induced to differentiate according to Buchholz (Buchholz et al., 2013). Basal media consisting of DMEM/F-12, N2 supplement, sodium pyruvate, pen/strep, NEAA, BSA (0.1%), HEPES (4mM), and NaHCO_3_ (0.1%) were used throughout the protocol, with additional components added as detailed below. On day 1, media were replaced with initiation media with Noggin (50ng/mL), DKK (10ng/mL), IGF (10ng/mL), nicotinamide (10mM) and Y-27632 (Rock inhibitor, 10μM). On day 3, intermediate media #1 containing Noggin (50ng/mL), DKK (10ng/mL), IGF (10ng/mL), bFGF (5ng/mL), and nicotinamide (10mM). On day 5, intermediate media #2 with DKK (10ng/mL), IGF (10ng/mL) and Activin A (100ng/mL) was used. From day 7 to 20, cells were kept in maturation media with Activin A (100ng/mL), SU5402 (10uM) and VIP (1uM). VIP was excluded from day 14 onwards, and cells were maintained in basal media until RPE emerges. RPE were either selected manually or entire wells trypsinized to expand the culture. RPE were then cultured in 5% RPE media consisting of MEMα, N1 supplement, FBS (5%), NEAA, pen/strep, hydrocortisone, triiodothyronine, taurine and Y-27632 (10μM). (Du et al., 2016; Sonoda et al., 2009) After 1 week in culture, culture media was changed to 1% FBS without Y-27632. All media components were purchased from Thermo Fisher Scientific (Waltham, MA, USA) or MilliporeSigma (Burlington, MA, USA), and exogenous components were purchased from PeproTech (Rocky Hill, NJ, USA).

Three iPSC clones were differentiated into RPE cell lines from each of the two SFD individuals (V-2 and V-7), and 1-5 RPE iPSC clones were differentiated into RPE cell lines from each of three normal iPSC age-matched controls (Supplementary Table 2). RPE between passages 2 to 6 were used, and cell lines were tested routinely for Mycoplasma contamination (MycoAlert, Lonza). Cells were passaged using 0.25% trypsin-EDTA and seeded on Matrigel® Matrix (Corning)-coated surfaces or on polyethylene terephthalate (PET) track-etched 0.4 μm pore size filters (Falcon) at minimum 2.0×10^5^ cells/cm^2^. Transepithelial resistance (TER) measurements were taken using Millcell ERS-2 epithelial volt meter (MilliporeSigma). RPE cells were cultured for 8 weeks before collection unless otherwise stated.

### 2.5. Genomic sequencing

DNA was purified from patient PBMCs for variant sequencing. Amplifications were carried out using Platinium® Taq DNA Polymerase High Fidelity kit (Life Technologies) with primers in Supplementary Table 3. Cycling conditions were 94° for 2 min followed by 35 cycles of 94° for 30 sec, 60° for 30 sec, 68° for 1 min, with a 2 min final extension at 68°. PCR products were purified with Edge Bio QuickStep 2 PCR purification (Edge BioSystems, Gaithersburg, MD), BigDye labeled using the BigDye Terminator v3.1 Cycle Sequencing Kit (Applied Biosystems, Foster City, CA) and sequenced with an ABI 3500 Genetic analyzer (Applied Biosystems).

### 2.6. CRISPR-Cas9 gene editing of the S204C TIMP3 mutation in SFD patient-derived iPSCs

iPSCs (1.0×10^6^ cells) from individual V-7 harboring the 610A>T (p.Ser204Cys) mutation in the TIMP3 locus were electroporated with Cas9 (0.15μM, Sigma) and guide (g)RNA (0.75μM, GGGGCCTATTTTCGTAGTAG, Synthego) as RNP complex along with ssDNA donor (4μM, IDT) using Amaxa nucleofector (Human Stem Cell kit 2) in presence of ROCK inhibitor and HDR enhancer (30μM, Integrated DNA Technologies, Coralville, IA, USA). Individual colonies were hand-picked and plated into 96 well plates. DNA was extracted using Quick Extract DNA extraction solution (Epicentre #QE09050) and nested PCR performed. The PCR product was purified using EXO-SAP enzyme (ThermoFisher) and sent for Sanger sequencing analysis. Gene edited iPSC colonies were expanded as described above.

To determine if the gRNA generated off-target effects from CRISPR-editing, we used CRISPOR (http://crispor.tefor.net/) to search for potential off-target sites and found 2 intronic regions with 3 mismatches. These regions (Off Target 1 Chr 6 88378477-499 and Off Target 2 Chr 22 33629462-484) were sequenced in the CRISPR-edited iPSC clones and aligned to the UCSB reference, which indicated no genetic alternations in these regions. A silent mutation was introduced in the PAM region and detected with sequencing, suggesting there were also no large deletions of the targeted gene (Supplementary Fig. 2).

### 2.7. Real-time PCR (qPCR)

iPSC RPE cultured were collected in TRIzol Reagent (Invitrogen) and RNA isolated with chloroform. RNA samples were purified treated with DNase (Roche) and RNeasy mini prep kit (Qiagen). cDNA was synthesized using iScript cDNA synthesis kit (Bio-Rad Laboratories, Hercules, CA, USA) and qPCR performed using Sybr Select Master Mix (Applied Biosystems) with primers in Supplementary Table 3. Relative gene expression was calculated after normalization to GAPDH and then to RPE control. Fold changes were represented as mean ± SD.

### 2.8. Western blotting

RPE were incubated in serum free medium for 48 hours before harvest. Media were collected for zymography/reverse zymography studies, and cells detached with 5mM EDTA were lysed in RIPA buffer with cOmplete™, Mini Protease Inhibitor Cocktail (Roche, Indianapolis, IN, USA). The culture plate was washed 6-7 times with 1xDPBS and once with ice cold sterile water to remove any remaining cells. ECM was scraped directly into Laemmli buffer (Bio-Rad). Bradford assay was used to determine protein concentrations. ECM and RPE cell lysate (15-20 μg protein) were run separately on precast Mini-Protean TGX gels (Bio-Rad) and transferred onto PVDF membrane (Immobilon). Blots were blocked with 5% BSA in PBS and incubated in ApoE (1:1000, AB947, Millipore), BEST1 (1:500, SAB2701026, Sigma-Aldrich), and RPE65 (1:5000, MAB5428, Millipore) overnight at 4°C. β-Actin (1:10,000, GTX629630, GeneTex) was used as a housekeeping gene. IRDye secondary antibodies were used and blots imaged with the Odyssey Clx (Li-Cor, Lincoln, NE, USA). For TIMP3 (1:1200, MAB3318, Chemicon), nitrocellulose membranes were used, while milk was used as blocking buffer and antibody diluent.

### 2.9. Zymography and reverse zymography

Zymography and reverse zymography were performed as previously described (Qi and Anand-Apte, 2015). In brief, media and ECM samples from 8-wk RPE were prepared in non-reducing Laemmli sample buffer and separated on either 7.5% SDS-polyacrylamide gel containing 1mg/ml gelatin (zymogram), or 12% SDS-polyacrylamide gels with 1mg/ml gelatin and RPE cell-conditioned media as a source of MMPs (reverse zymography). After electrophoresis, gels were incubated in a solution of 25mg/ml Triton X-100 to renature proteins. The gels were then washed with water and incubated 16 hours in 50mM Tris-HCL (pH7.5) containing 5mM CaCl_2_ and 0.2mg/ml sodium azide at 37°C. Gels were then stained with 5mg/ml Coomassie Blue R-250 in acetic acid/methanol/water (1:3:6) for 2 hours and destained with acetic acid/methanol/water (1:3:6).

### 2.10. Transmission electron microscopy

iPSC-derived RPE cells on PET-filters were fixed in 4% glutaraldehyde in 0.1M cacodylate buffer. Filter membranes were excised from the plastic culture inserts, stained with 1% tetroxide for 1hour, uranyl acetate solution for 20min, and rinsed before a 30%-100% ethanol dehydration series. Dehydrated samples were embedded in Epon Araldite 502 (Electron Microscopy Sciences, Washington, PA, USA) before curing for 24 hours at 60°C. 80nm sections were imaged on a JEOL JEM 1200EXII electron microscope at 80kV and a spot size of three. For previously paraffin-embedded SFD globes, approximately 1mm x1mm-sized pieces were de-paraffinized through three consecutive washes in xylene followed by graded ethanol dilutions and into PBS. These were then fixed for TEM as described above. TEM images were montaged into a panorama view with an average of 27 images per montage, ranging between 91,517.926nm and 126,419.400nm in length. For quantitative analysis, deposits with at least two of the characteristic structures of basal disruption, lipid droplets, banded collagen, and/or electron dense debris were counted. Extracellular matrix area and sub RPE deposit dimensions were measured using FIJI software (ImageJ, National Institutes of Health, Bethesda, MD, USA), and ECM area measurements were adjusted for montage length to determine average ECM thickness with standard error. ECM measurements from SFD sections did not include deposits. Six control and six SFD montages were quantified.

### 2.11. Immunostaining

Paraffin sectioned from adult SFD and age-matched control globes were de-paraffinized and antigen retrieval performed using 10mM sodium citrate buffer with 0.05% Tween-20. Tissues were blocked with 10% horse serum and incubated overnight with primary antibodies at 4°C. Antibodies included ApoE (1:1000, AB947), Bestrophin (1:250, MAB5466, Millipore), Collagen VI (1:250, ab6588, Abcam, Cambridge, MA), Clusterin (1:200, AB825, Millipore), CRALBP (1:1000, gift from Dr. Jack Saari, UW Vision Core, Seattle, WA), TIMP2 (1:500, H00007077-M03J, Novus Biologicals, Centennial, CO), TIMP3 (1:5000, GTX25939, GeneTex, Irvine, CA), and Vitronectin (1:1000, AB19014, Millipore). Alexa Fluor secondary antibody (Molecular Probes, Grand Island, NY) were used at 1:500 dilution. Sections were counterstained with DAPI (Sigma-Aldrich) and mounted with Fluoromount-G (SouthernBiotech, Birmingham, AL, USA). Images were taken at 63x using Leica DM6000 CS confocal microscope (Leica Microsystems). For iPSC-RPE cells, filter inserts were fixed in 4% PFA, entered a sucrose gradient, excised from the plastic holder and embedded in Tissue-TEK OCT compound (Sakura Finetek USA, Inc, Torrance, VA, USA). Blocks were cut on the Leica CM1850 cryostat to obtain 20 μm thick sections. Staining was performed as described above.

### 2.12. LDH assay and ethidium homodimer staining

RPE cells in 96-well plate were treated with 1mM H_2_O_2_ in phenol-free DMEM supplemented with 5.5 mM glucose, 1% FBS and pen/strep. A 20μL aliquot of media was removed at 48 and 72 hours for LDH assay per manufacturer’s protocol (Genzyme, Cambridge, MA, USA). After 72 hour treatment, culture media was replaced with ethidium homodimer stain (Invitrogen) diluted in Kreb’s Ringer buffer (KRB). Following 10min incubation, images were taken on the EVOS microscope (AMG, Fisher Scientific) and red fluorescent cells were counted from each well.

### 2.13. Targeted metabolomics analysis with liquid chromatography and mass spectrometry analysis

To collect cells for metabolomics, RPE cells were rinsed once with cold 154mM NaCl solution, and scraped in cold 80% methanol on dry ice. The metabolites were separated with an ACQUITY UPLC BEH Amide analytic column (2.1 X 50 mm, 1.7μm, Waters) by a Shimadzu LC Nexera X2 UHPLC coupled with a QTRAP 5500 LC MS/MS (AB Sciex). Each metabolite was tuned with standards for optimal transitions (Supplementary Table 4). The extracted multiple reaction monitoring peaks were integrated using MultiQuant 3.0.2 software.

### 2.14. Stable isotope-labeled metabolite analysis

RPE media were replaced with DMEM supplemented with FBS (1%), glucose (5.5mM) and ^13^C glutamine (200 μM, Sigma) or ^13^C proline (200 μM, Sigma) for 24 hours. Metabolites were extracted in 80% cold methanol, dried, derivatized by methoxyamine and N-tertbutyldimethylsilyl-N-methyltrifluoroacetamide, and analyzed by an Agilent 7890B/5977B GC/MS system with an Agilent DB-5MS column (30 m × 0.25 mm × 0.25 μm film). Mass spectra were collected from m/z 80–600 under selective ion monitoring mode. The data was analyzed by Agilent MassHunter Quantitative Analysis Software.

### 2.15. Alizarin staining

RPE cells were cultured on 8-well chambered Permanox slides (Nunc Lab-Tek, ThermoFisher Scientific). Cells were fixed in 4% PFA and stained with Alizarin Red (pH 6.4) according to standard protocol. Brightfield images were taken on a Nikon Eclipse E1000 wide-field microscope and Alizarin positive area quantified on FIJI. Staining/imaging were performed on RPE derived from two normal control iPSC lines, two SFD iPSC lines, and two CRISPR-corrected SFD clones (n=7-12 wells for each group). For specific cell line information, see Supplementary Table 2.

### 2.16. Statistical Analysis

Data were analyzed in GraphPad Prism (GraphPad Software, La Jolla California USA) using unpaired two-tailed t-tests or one-way ANOVA. Significance was determined by p-value <0.05 (*), <0.01(**), and <0.001 (***). Data presented as mean ± SD unless otherwise stated. Volcano plot was made and analyzed with XLSTAT (2018.1.49540) in Microsoft Excel using unpaired t-test.

## 3. Results

### 3.1. Sorsby Fundus Dystrophy (S204C TIMP3 variant) family members demonstrate typical clinical and pathologic features of the disease

Members of a large Sorsby Fundus Dystrophy (SFD) family underwent sequencing of exon 5 in the *TIMP3* gene for the S204C (c.610A>T) variant and results are included in an updated pedigree (Fig. 1A) (Carrero-Valenzuela et al., 1996). Induced pluripotent stem cells (iPSCs) were generated from two affected individuals, V-2 and V-7, with different disease severity (Fig. 1B). Individual V-2, a 66-year old woman, experienced loss of central vision in both eyes in her fourth decade as a result of complications from choroidal neovascularization (CNVM). Her best corrected visual acuity (BCVA) was 1/200 in the right eye and 2/200 in the left eye. There was widespread RPE and choroidal atrophy with central macular pigmentary clumping and disciform scarring, and OCT imaging revealed extensive subretinal scarring with severe overlying macular atrophy (Fig. 1B). Individual V-7, a 63-year old woman, was asymptomatic upon initial presentation with 20/20 BCVA in both eyes. Although she developed CNVM in the right eye several years later, it was successfully with a single injection of intravitreal bevacizumab. Individual V-7 has RPE mottling and large drusen in the macula, along the vascular arcades and mid-periphery. OCT imaging shows both conventional drusen (Fig. 1B, white arrow) and subretinal drusenoid deposits (SDD), with the typical peaked morphology (Fig. 1B, red arrowhead) (Gliem et al., 2015).

**Figure 1.**
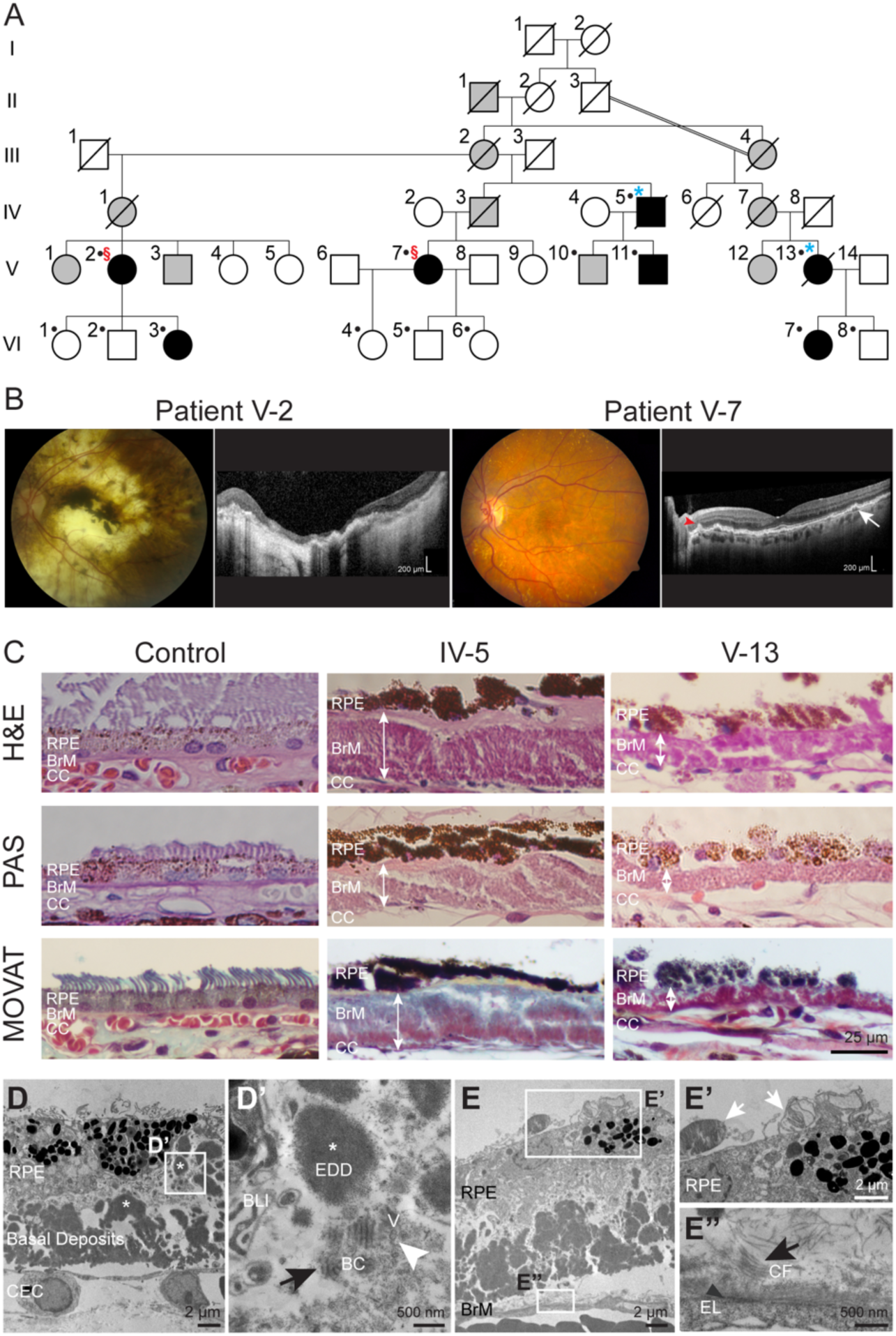
Clinical imaging and histological findings of members of a S204C *TIMP3* Sorsby Fundus Dystrophy family. (A) Abridged pedigree of SFD family. Filled black symbols indicate individuals with SFD determined by clinical examination. Slashed symbols indicate deceased individuals. Dots indicate genetic confirmation of disease status. §, individuals from whom iPSCs were generated; *, donated globes. (B) Retinal imaging of the left eyes of patient V-2 revealed extensive RPE and choroidal atrophy with subretinal scarring. Patient V-7 had macular RPE pigment mottling and extensive yellow deposits. Drusen (white arrow) and conical-shaped subretinal drusenoid deposits (red arrowhead) were seen with OCT imaging. (C) Globes from SFD family members IV-5 and V-13 had marked deposition of sub-RPE material (white double arrowheads) internal to the elastin layer of the BrM. Movat’s stain demonstrated the presence of mucopolysaccharides (blue-green), collagen (yellow) and a thickened and disrupted elastin (black) sublayer of BrM. (D) TEM imaging shows RPE with irregular morphology overlying a thick layer of electron dense deposits (EDD, white asterisk). (D’) Wide-spaced banded collagen (BC, black arrow) of 110nm periodicity, and vesicular structures (V, white arrowhead) were also seen amongst the deposits. (E’) Partial phagocytosis of photoreceptor outer segments (white arrows) by RPE. (E’’) Disorganized BrM with abundant collagen fibrils (CF, black arrow) are noted above the residual discontinuous elastic layer (EL, black arrowhead). BLI – basal laminar infoldings; CEC – choroidal endothelial cell.

Histology was performed on donated globes from two genetically confirmed family members, IV-5 and V-13, ages 85 and 77 years, respectively, at the time of globe donation. Light microscopy showed disruption of the RPE monolayer with either missing or multiple layers of RPE, and the RPE cells themselves exhibited irregular morphology (Fig. 1C). As is pathognomonic to SFD, both globes were observed to have thick (10-50 μ ) layers of sub-RPE deposits, which were located internal to the elastin layer of Bruch’s membrane. The deposits contained mucopolysaccharides and collagen as shown by Movat’s pentachrome staining.

Transmission electron microscopy (TEM) of the sub-RPE deposits in SFD globes showed large regions of electron dense deposits, wide-spaced banded material, and electron-lucent spaces of varying diameter (Fig. 1D). Presumed to be type VI collagen, wide-spaced material with ∼80-110 nm periodicity has been shown in SFD globes (Chong et al., 2000; Knupp et al., 2002). BrM was severely disrupted throughout (Fig. 1E), and RPE cells had abnormal morphology. Rare phagocytosis of outer segments was noted (Fig. 1E’, white arrows), implying some remaining RPE function. Disorganized BrM with abundant collagen fibrils were noted above the residual discontinuous elastic layer (Fig. 1E’”).

### 3.2. Sorsby Fundus Dystrophy iPSC-derived RPE have high levels of intracellular and ECM TIMP3 protein while MMP-2 and -9 activity is unchanged

Multiple iPSC clones were generated from three controls and two SFD individuals (V-2 and V-7) and differentiated into RPE (Supplementary Table 2). By four weeks in culture, SFD patient and control iPSC-derived RPE cell lines demonstrated typical characteristic features of mature RPE, including hexagonal morphology, tight junction formation, and the presence of melanin pigment granules (Fig. 2A). RPE cell lines seeded on filter inserts established tight junctions and transepithelial resistance (TER) measurements of ≥ 200Ω•cm2 (Fig. 2B).

**Figure 2.**
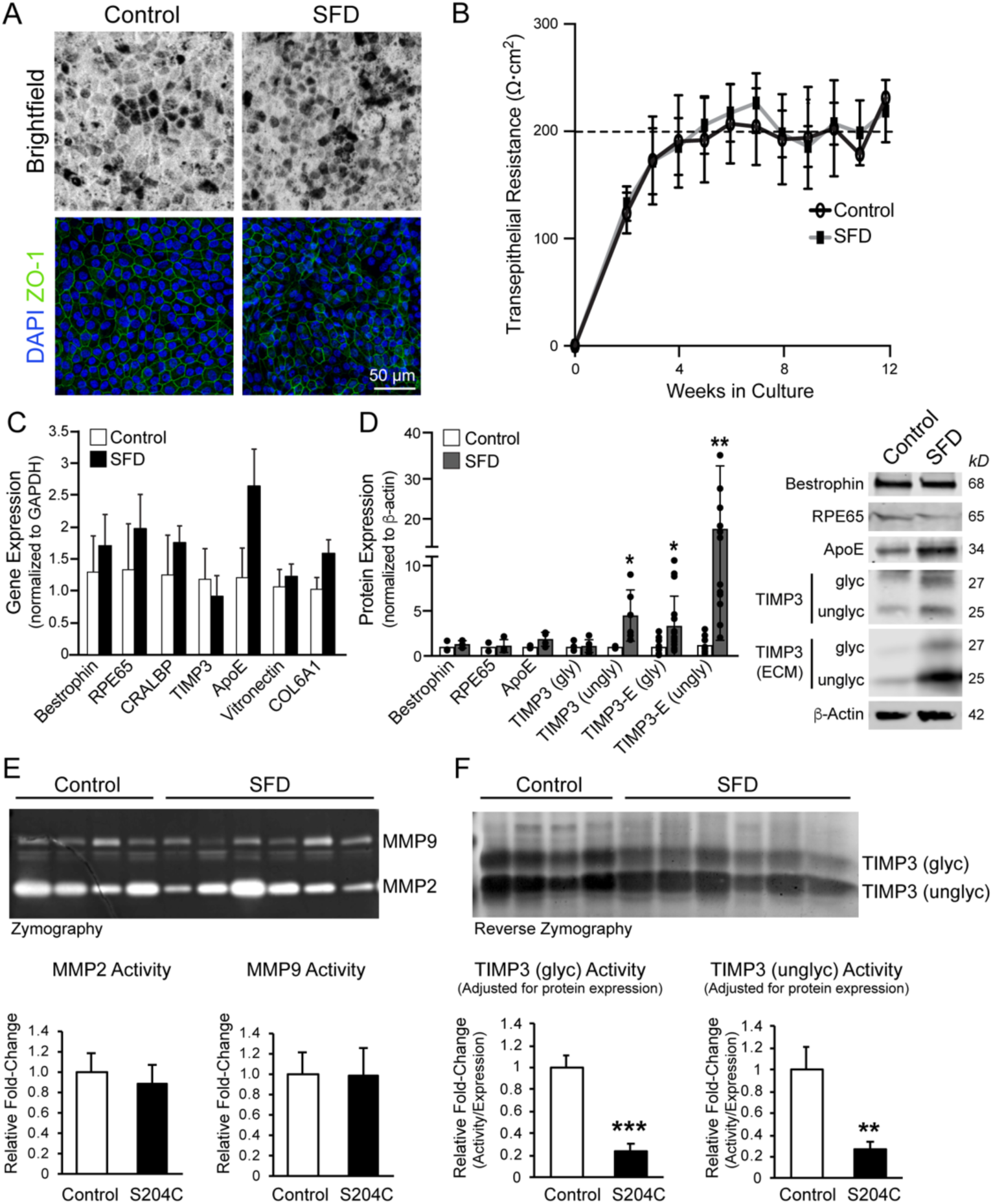
TIMP3 protein expression and activity in SFD RPE. (A) Brightfield images and ZO-1 staining showed typical RPE pigmentation, hexagonal morphology, and presence of tight junctions in both control and SFD iPSC RPE cells. (B) Both control and SFD RPE demonstrated TER of ≥200 Ω •cm^2^ by 5-6 weeks in culture. (C) Transcript levels of RPE-related genes were unchanged between control and SFD RPE, although (D) intracellular unglycoslyated TIMP3 protein and unglycoslyated TIMP3 in the ECM both increased ∼4-fold and ∼18-fold, respectively. Expression of intracellular glycosylated TIMP3 was similar between SFD and control RPE lines, although it was increased in the ECM of SFD RPE. (E) Zymography with cell culture media showed no difference in either MMP-2 or MMP-9 activity in SFD RPE compared to controls, where hydrolysis of the gelatin substrate by proteinases is reflected as white bands on a dark background. (F) In reverse zymography where TIMP3 inhibition of MMP activity is represented by dark bands against a lighter background, no significant difference was seen in overall TIMP3 activity between SFD and control RPE, although both glycoslylated and unglycosylated TIMP3 activity in SFD RPE was ∼5-fold decreased when adjusted for TIMP3 expression. Mean ± SD; *p<0.05; **p<0.01, ***p<0.001; n=3.

Although there is a wide range of TER measurements reported in the literature, RPE cells grown on filter inserts with plating substrates similar to ours use TER ≥ 200Ω•cm^2^ as an indication of cellular maturity and polarization (Brydon et al., 2019; Galloway et al., 2017; Gong et al., 2020; Hazim et al., 2017; Jin et al., 2002; Kuroda et al., 2019).

There was no significant difference in transcript expression of bestrophin, RPE65, CRALBP, vitronectin, and COL6A1 in SFD-derived RPE compared to controls (Fig. 2C). Although APOE mRNA expression trended higher in SFD RPE, it did not reach statistical significance. There was no change in TIMP3 mRNA expression in SFD-derived RPE, although SFD iPSC RPE showed ∼4-fold higher intracellular unglycosylated TIMP3 protein expression and ∼18-fold higher TIMP3 deposition in the ECM compared to controls (Fig. 2D). These findings are consistent with recently reported findings from similar SFD iPSC RPE lines (Hongisto et al., 2020), although we find that glycosylated TIMP3 was also increased in the ECM of SFD RPE. These findings reproduce *in situ* hybridization results in SFD globes that demonstrate no differences in levels of TIMP3 transcript despite significant TIMP3 protein deposition (Bailey et al., 2001; Chong et al., 2000; Chong et al., 2003; Fariss et al., 1998).

Zymography was used to assess the activity of two matrix metalloproteinases (MMPs) commonly found in Bruch’s membrane, MMP-2 and MMP-9. Hydrolysis of the gelatin substrate by proteinases is reflected as white bands on a dark background. We found no significant difference in MMP-2 or MMP-9 activity in SFD RPE compared to controls (Fig. 2E). We then examined the inhibitory activity of mutant TIMP3 from SFD cell lysates using reverse zymography (Fig. 2F). In reverse zymography, TIMP3 inhibition of MMP activity is represented by dark bands against a lighter background of digested gelatin (Qi et al., 2009). We observed a small decrease in MMP inhibition from SFD RPE (Fig. 2F, top), although when adjusted for levels of TIMP3 protein, SFD TIMP3 displayed a ∼5-fold decrease in MMP inhibition compared to controls (Fig 2F, bottom). These results show that while there is significantly greater intracellular and ECM TIMP3 in SFD RPE, mutant TIMP3 is 5-fold less effective at inhibiting MMP-2 and MMP-9.

### 3.3. S204C TIMP3 iPSC-derived RPE form thinner extracellular matrices and elaborate large deposits similar in composition to basal laminar drusen

We next examined the ultrastructure of SFD RPE cultured on polyethylene terephthalate (PET) culture inserts by TEM. Control iPSC RPE cell lines consistently deposited a uniform layer of fine filaments at the filter interface, which on average measured ∼800 nm in thickness (Fig. 3A, top image, arrow). In contrast, the layer of filamentous ECM elaborated by SFD RPE was irregular and significantly thinner (Figs. 3B). Quantification of ECM thickness measurements show a 4-fold decrease in SFD RPE ECM thickness compared to that of controls (Fig. 3C). At 8 weeks in culture, SFD-derived RPE elaborated large accumulations of extracellular deposits while no deposits were detected in control RPE at the same time point (Fig. 3D). SFD RPE overlying the large deposits maintained apical microvilli, basal infoldings and tight junctions (Fig. 3E). The deposits contained short segment, long spaced transverse bands of 30nm in width and ∼110 nm axially repeated periodicity, consistent to what was recently described in deposits of other SFD iPSC RPE lines (Hongisto et al., 2020). These complexes were similar in size and architecture to the wide-spaced banded structures seen in sub-RPE deposits of their family member SFD globes (Fig. 1D’). Multiple electron lucent vesicles of varying diameter, consistent with lipid, were also noted in the culture deposits (Fig. 3E’”, white arrowheads), which were similar to similar vesicles detected in SFD globes (Fig. 1D, white arrowhead). While large electron dense accumulations were observed in SFD globes (Fig. 1D), neither the SFD iPSC RPE deposits in our study nor those previously reported (Hongisto et al., 2020) reproduced these accumulations in culture.

**Figure 3.**
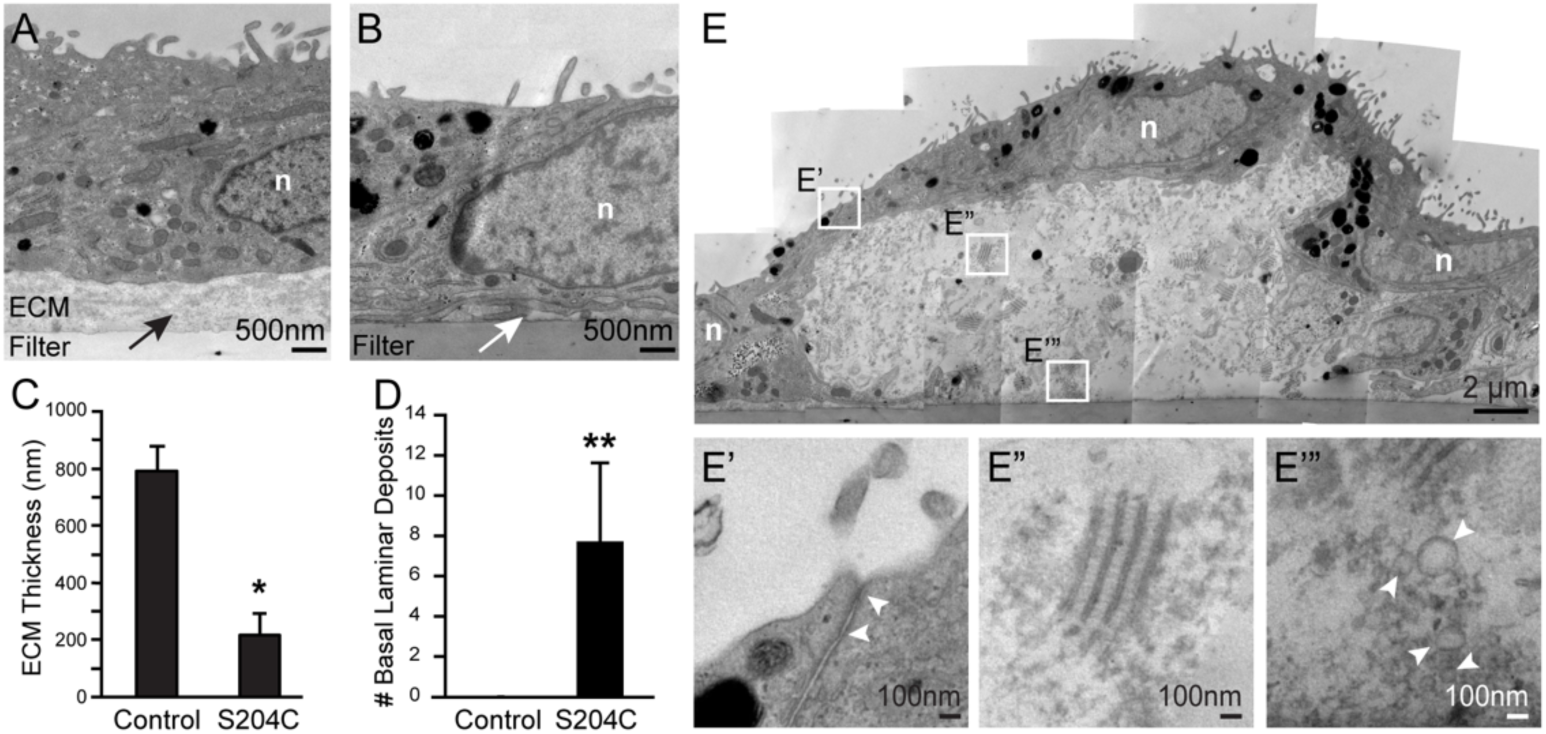
SFD iPSC-derived RPE form irregular extracellular matrices and large basal deposits in culture. (A) Control RPE cultured for 8 weeks on PET filter elaborated a uniform layer of ECM (black arrow) while (B) SFD RPE formed irregular and markedly thinner filamentous ECM (white arrow). (C) SFD RPE ECM was significantly thinner (∼4-fold) while also having (D) significantly increased number of basal laminar deposits. (E) One representative sub-RPE deposit measures 15μm in length and 6μm in height. Overlying RPE cells maintain typical ultrastructural features including apical microvilli, basal infoldings and tight junctions (E’). Sub-RPE deposits contained numerous short segmented, long-spaced transverse bands of 30nm in width and ∼110 nm axially repeated periodicity (E’’), consistent with banded collagen seen in human globes. (E’’’) Electron lucent vesicles of varying diameter, possibly representing lipid particles, were also noted in the deposits (white arrowheads). n, nucleus. Mean ± SD; **p<0.01; n=3 per group.

### 3.4. SFD iPSC-RPE deposits are similar in composition to sub-RPE deposits in SFD globes

Sub-RPE deposit components of iPSC RPE cultured on filter membranes were compared with SFD globe deposits by immunohistochemistry. RPE express TIMP3 and secrete it into BrM, where it is tightly bound to the extracellular matrix. Increased levels of TIMP3 in BrM are found in aged eyes and even higher levels have been noted in SFD patients globes (Chong et al., 2000; Fariss et al., 1998). Immunostaining of control globes demonstrated TIMP3 in RPE and less prominently in BrM (Supplementary Fig. 3A), whereas in SFD globes, there was significant TIMP3 staining in the sub-RPE deposits (Fig. 4A). There was strong TIMP3 immunoreactivity in the sub-RPE deposits of SFD iPSC RPE (Fig. 4B). Immunostaining of other basal laminar drusen components including apolipoprotein E (ApoE), vitronectin, clusterin (Fig. 4C-D), and collagen type VI (Supplementary Fig. 3G-H) was also performed and showed similar levels of immunoreactivity between iPSC RPE and SFD globe deposits. TIMP2, a non-secreted intracellular RPE protein, was detected in RPE cells but not in the deposits of SFD iPSC RPE or SFD globes (Fig. 4E-F). Overall the pattern and degree of staining in SFD globes and SFD iPSC RPE deposits are similar among the basal laminar drusen components tested.

**Figure 4.**
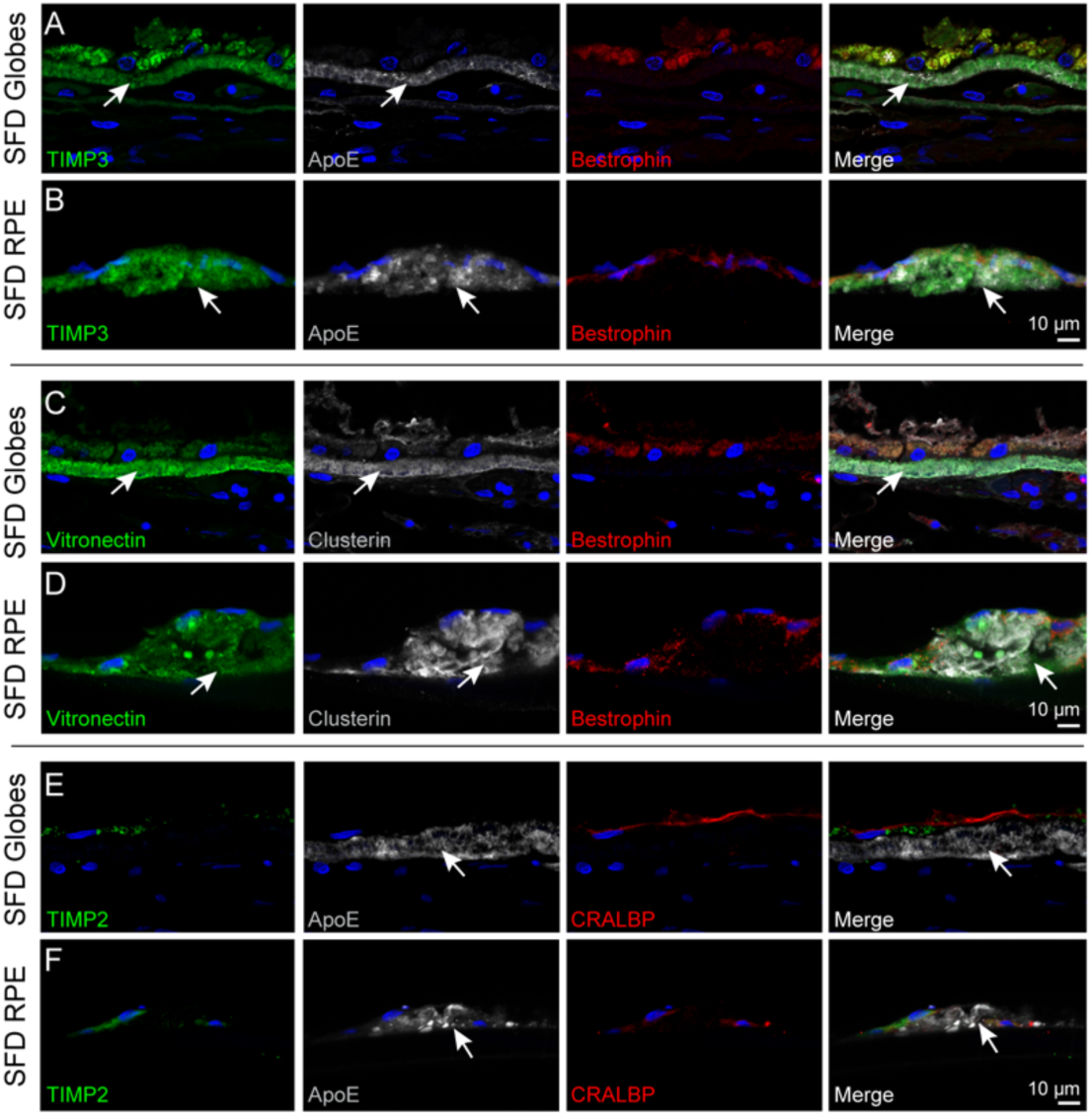
SFD iPSC-RPE deposits in culture are similar in composition to sub-RPE deposits seen in SFD patient globes. (A) In SFD globes, TIMP3 was observed in the RPE and in the thickened BrM. ApoE, a major component of drusen, was also found in the BrM. (B) Deposits formed by the bestrophin-positive SFD iPSC RPE similarly immunostained with TIMP3 and ApoE, and the signals did not appear to colocalize. (C,D) Vitronectin and clusterin staining were localized to the basal deposits in both SFD globes and iPSC-RPE cells, indicating similarities in composition between the globe and *in vitro* model. (E,F) As expected, TIMP2, a non-secreted intracellular protein, was detected only in the RPE of SFD globes and iPSC RPE, and was not detected in the ApoE-positive deposits.

### 3.5. S204C TIMP3 iPSC-derived RPE have increased intracellular 4-hypdroxyproline and are more susceptible to oxidative stress

Because cellular metabolism can be affected by disturbances in ECM homeostasis, we examined whether the metabolism of SFD iPSC RPE is altered compared to control RPE. Forty-six key metabolites involved in energy metabolism including glycolysis, mitochondrial Krebs cycle, amino acids, ATP and NADH were examined using targeted metabolomics with LC MS/MS (Supplementary Table 4). 4-hydroxyproline is the most significantly altered of the metabolites with a ∼2.7-fold increase and p-value ≤ 1.0 x 10 (Fig. 5A). 4-hydroxyproline is a major component of collagen and required for collagen stability (Gordon and Hahn, 2010). Its increase could result from either greater synthesis from proline hydroxylation or increased degradation from collagen. To determine if the increase in 4-hydroxyproline was due to increased synthesis, we used a labeled proline tracer. We incubated the RPE cells with labeled ^13^C proline but did not detect a significant difference in the consumption of ^13^C proline in the cell culture media. This suggests that increased 4-hydroxyproline might not result from *de novo* synthesis but from degradation (Fig. 5B).

**Figure 5:**
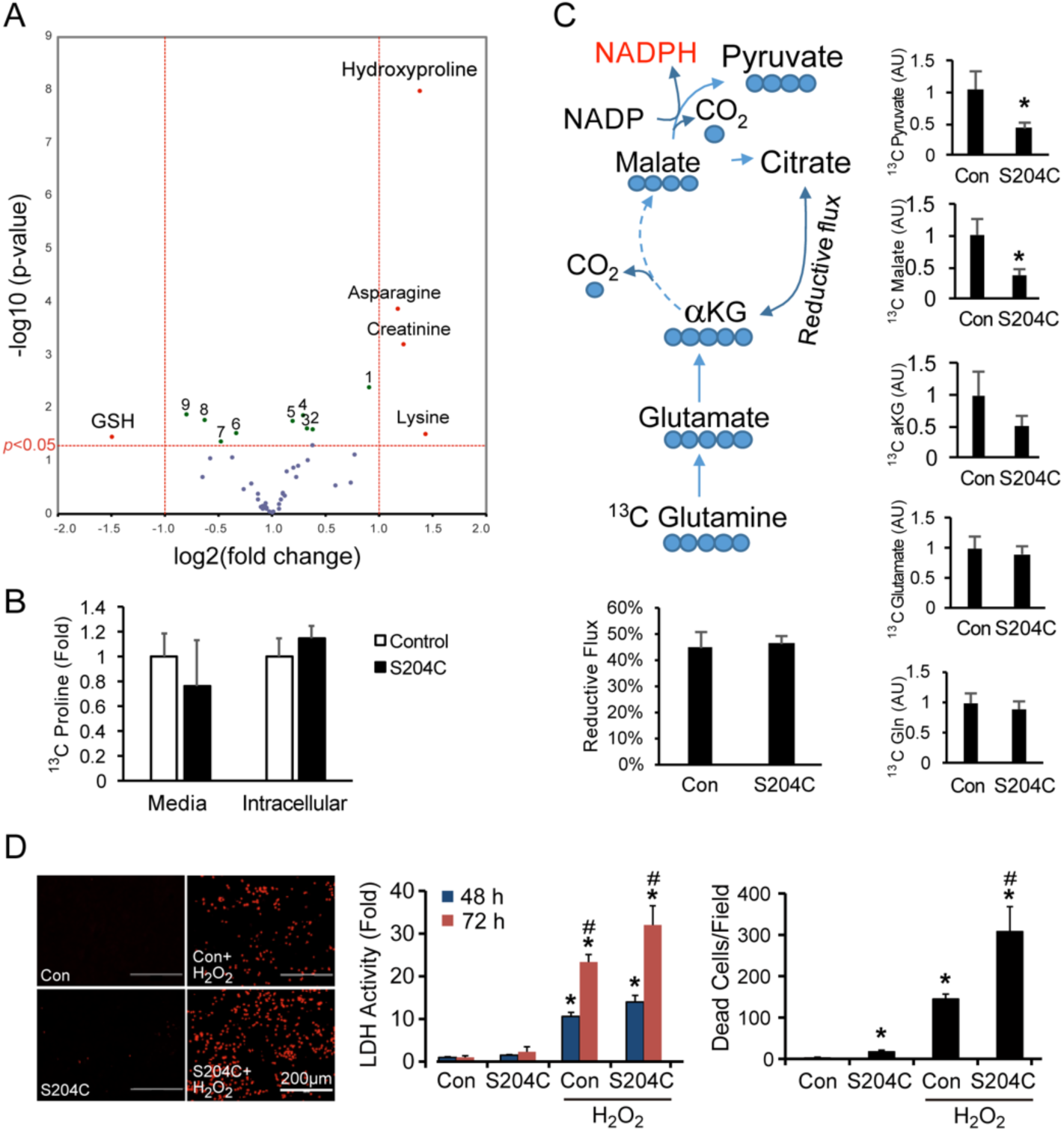
SFD iPSC-derived RPE have altered metabolism and are more susceptible to oxidative stress. (A) Metabolite analysis of SFD RPE represented in a volcano plot showed ∼2.7-fold higher levels of hydroxyproline (major product of collagen degradation), and ∼3-fold reduced glutathione (GSH) levels. Significant changes in other metabolites include (1) tyrosine, (2) aspartate, (3) alanine, (4) glutamate, (5) fumarate, (6) GTP, (7) FAD, (8) GSSG, (9) choline. (B) SFD RPE incubated with labeled ^13^C proline did not result in a significant increase in ^13^C proline consumption, suggesting that increased 4-hydroxyproline levels were unlikely due to increased synthesis. (C) Reductive carboxylation was examined by quantifying ^13^C labeled metabolites from RPE incubated with ^13^C glutamine. There was no change in reductive flux, although the flux to malate and pyruvate was significantly attenuated in SFD RPE. (D) SFD RPE treated with 1mM H_2_O_2_ for 48 or 72 hours had elevated LDH activity and cell death compared to their control counterparts. Mean ± SD; *p<0.05 vs control without treatment; ^#^p<0.05 vs con with H_2_O_2_.; *n* = 4.

Glutathione in its reduced state (GSH) is a metabolite that is critical in preventing damage from oxidative stress, and we found that it is significantly decreased by ∼3-fold in SFD iPSC RPE compared to controls (Fig. 5A), indicating that SFD RPE may have higher levels of oxidative stress. This finding is consistent with a recent report indicating that TIMP3 mutations may result in dysregulation of antioxidant homeostasis in a SFD mouse model (Wolk et al., 2020). We previously reported that RPE has an exceptionally high capacity for reductive carboxylation and that reductive carboxylation can play a significant role in RPE resistance to oxidative stress (Du et al., 2016). We traced reductive carboxylation by quantifying ^13^C labeled metabolites from SFD iPSC RPE and control iPSC RPE incubated with ^13^C glutamine for 24 hours. The reductive flux in the iPSC RPE cells is as high as mature human primary RPE cells; however, there is no difference in reductive carboxylation between SFD and control iPSC RPE (Fig. 5C). Moreover, SFD RPE cells produce lower levels of ^13^C-labeled malate and pyruvate but not α ketoglutarate (Fig 5C). The production of pyruvate through malate is an important source of NADPH which is required for GSH regeneration, indicating that SFD RPE may be more oxidatively stressed at baseline. We did not detect significant changes in the levels of lactate, pyruvate, pentose phosphate pathway intermediates or glucose consumption (Fig. 5A). To determine whether SFD iPSC RPE cells are more sensitive to oxidative damage, SFD and control iPSC RPE were treated with 1mM H_2_O_2_, a strong oxidative stressor. After 48 hours, there were more than twice as many dead cells in the SFD iPSC RPE than in control RPE as detected by ethidium homodimer staining (Fig. 5D). Consistently, LDH activity was also highest in the culture media from SFD iPSC RPE cells treated with H_2_O_2_ (Fig. 5D). Together, these data indicate that SFD iPSC RPE are more oxidatively stressed at baseline and more vulnerable to oxidative damage.

### 3.6. Calcium deposits are found in SFD Globes and SFD iPSC-RPE, while CRISPR-edited iPSC RPE form significantly fewer deposits

Calcium aggregates are found in refractile drusen and in the elastic layer of aged BrM (Davis et al., 1981; Hogan et al., 1971; Suzuki et al., 2015). Hydroxyapatite spherules were identified in sub-RPE deposits of older eyes, and porcine RPE form deposits containing hydroxyapatite after aging in culture (Pilgrim et al., 2017; Thompson et al., 2015). Alizarin red, an anthroquinone derivative, identifies free calcium and calcium phosphates, in tissues sections and cultured cells. While there was little Alizarin red staining in normal globes, diffuse staining of the thickened BrM deposits and underlying enlarged choriocapillaris of SFD eyes was observed (Fig. 6A). SFD iPSC RPE cultured on chambered slides formed abundant calcium deposits, whereas control RPE formed significantly fewer (Fig. 6B). CRISPR gene editing was performed on one SFD patient’s iPSC clone to correct the S204C TIMP3 mutation, and two corrected iPSC clones (cSFD) were differentiated into RPE (Supplementary Fig. 2). CRISPR-corrected TIMP3 SFD RPE showed hexagonal morphology, formed tight junctions, and formed significantly fewer basal deposits (Fig. 6D-E). Glycosylated TIMP3 expression was not significantly decreased in CRISPR-edited SFD RPE compared to uncorrected SFD RPE, although unglycosylated TIMP3 was significantly decreased in CRISPR-edited RPE, albeit at a higher level than control iPSC RPE (Fig. 6F-G). CRISPR-editing of SFD RPE resulted in significantly fewer calcium-containing deposits compared to SFD RPE (Fig. 6B-C).

**Figure 6.**
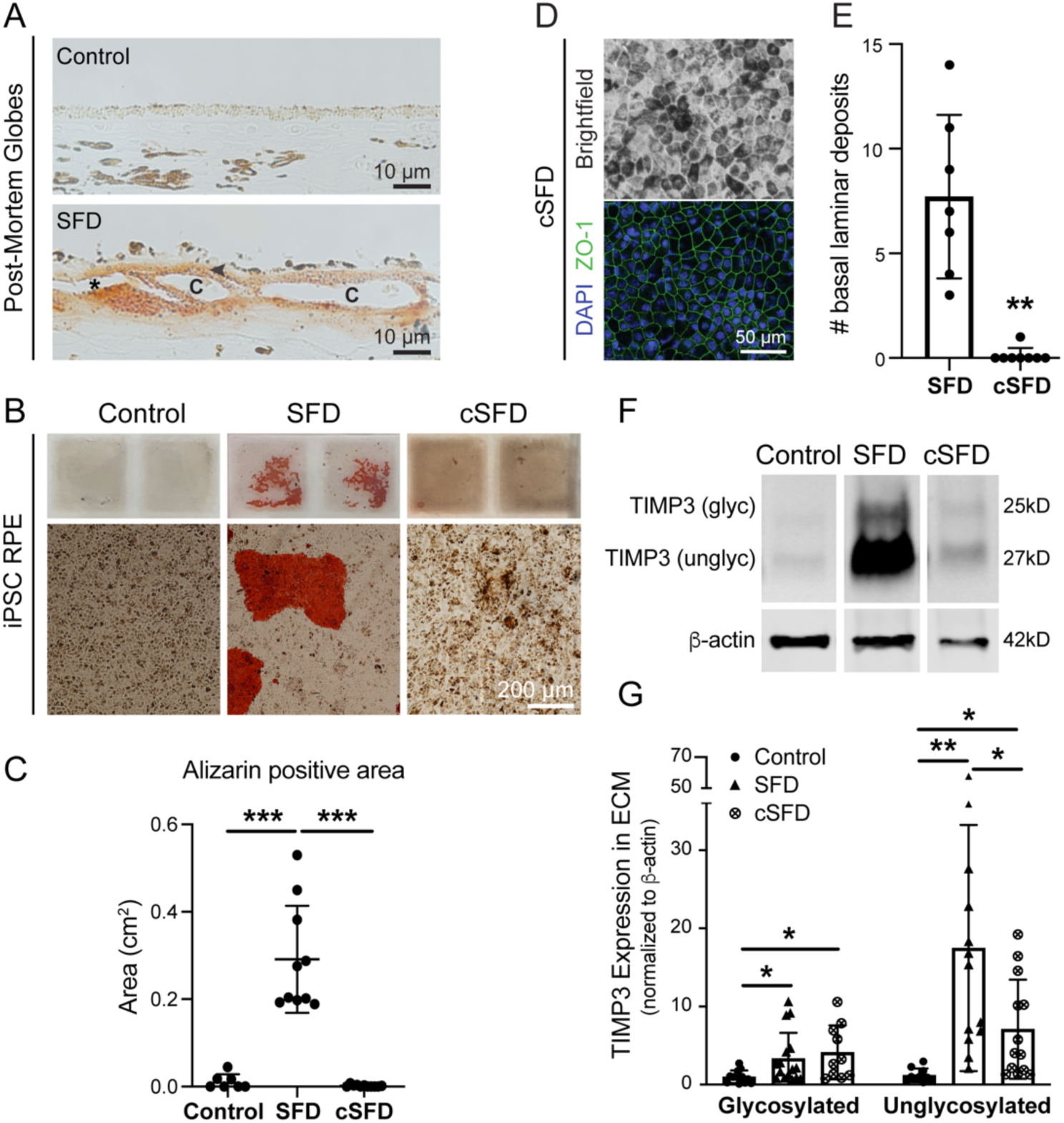
CRISPR-edited iPSC RPE (cSFD) form fewer deposits and have lower unglycosylated TIMP3 expression. (A) Globe sections stained with Alizarin red for calcium showed staining in thickened deposits basal to RPE cells (arrowhead) and around the choriocapillaris (c). *artifactual separation. (B) SFD iPSC RPE cultured on slides form large sub-RPE deposits that stained intensely with Alizarin red. Control and cSFD RPE show similarly low levels of calcium staining. (C) Quantification of Alizarin-positive areas. *n* = 7-12/cell type (D) cSFD RPE have typical pigmentation, hexagonal morphology and ZO-1 staining. (E) cSFD RPE have significantly fewer deposits by TEM imaging compared to uncorrected SFD RPE at 8 wks in culture. (F,G) cSFD has significantly decreased unglycosylated TIMP3 in the ECM compared to uncorrected SFD RPE cells, although both glycosylated and unglycosylated TIMP3 levels remained higher compared to normal control iPSC RPE. *p<0.05; **p<0.01; ***p<0.001

## 4. Discussion

A pathognomonic feature of Sorsby fundus dystrophy globes is a thick layer of extracellular deposits between the basal lamina of the RPE and the inner collagenous layer of Bruch’s membrane (BrM) (Chong et al., 2000; Fariss et al., 1998; Kuntz et al., 1996). We similarly found that SFD family member globes with the S204C *TIMP3* mutation in our study had a ∼25μm thick layer of sub-RPE deposits located internal to a disrupted elastin layer of Bruch’s membrane (BrM). We found that the deposits were composed of collagen, mucopolysaccharides, and other components of basal laminar drusen (BlamD) (Curcio, 2012; Johnson et al., 2011; Sarks et al., 2007; Wang et al., 2010), including apolipoprotein E, vitronectin, clusterin, and TIMP3. BlamD are also found in aging retinas, albeit to a much lesser extent than in SFD, and there is a strong relationship between macular BlamD and the formation of exudative age-related macular degeneration (AMD) (Khan et al., 2016). The accumulation of BlamD causes the RPE to slowly separate from the underlying BrM, possibly decreasing nutrient transport and RPE function (Jacobson et al., 1995; Zhang et al., 2019).

SFD iPSC-derived RPE lines generated from two patients formed significantly more sub-RPE deposits (∼6-90 μm in height) compared to control RPE lines at 8 weeks (∼60 days) in culture. While these deposits were similar in composition to BlamD found in SFD family member globes as observed by immunofluorescence staining, TEM imaging revealed additional similarities, including the presence of long-spaced collagen, presumed to be collagen type VI (Knupp et al., 2002), and lipids. TEM images further revealed that in areas without deposits, SFD RPE had decreased ECM thickness compared to controls, suggesting that SFD RPE may have dysregulated ECM homeostasis in addition to their propensity to forming large basal deposits. It is notable that prior studies differ on whether SFD iPSC RPE accumulate significantly more deposits than control RPE lines (Galloway et al., 2017; Hongisto et al., 2020). Possible factors contributing to differences in these studies include patient-specific non-*TIMP3* genetic modifiers, length of time in culture, and plating substrates (e.g. Matrigel, collagen, laminin). However, under our culture conditions, when CRISPR-Cas9 gene editing was performed to correct the S204C mutation in one SFD patient’s iPSCs, the number of basal deposits formed on filter inserts by the CRISPR-corrected SFD iPSC-derived RPE lines was significantly reduced compared to uncorrected SFD RPE and was similar to that of control RPE. Further, TIMP3 gene correction appeared sufficient to reverse the prominent deposition of free calcium and calcium phosphates exhibited by SFD iPSC RPE. In our SFD family member globes, we detected calcium deposits in both BrM and the enlarged choriocapillaris, which is consistent with calcium deposits found in a previously described S204C TIMP3 SFD donor eye (Chong et al., 2000). Although specific mechanisms linking the S204C TIMP3 mutation to extracellular calcium deposition is unclear, the complex array of sub-RPE calcium deposits observed in AMD patients has been proposed to result in part from the release of mitochondrial calcium in failing RPE (Pilgrim et al., 2017; Tan et al., 2018; Thompson et al., 2015).

Increased TIMP3 protein expression by SFD iPSC RPE was not accompanied by an increase in TIMP3 mRNA expression, which is consistent with *in vivo* findings in SFD globes where high levels of TIMP3 protein in BlamD were not accompanied by an increase in RPE TIMP3 mRNA expression (Bailey et al., 2001; Chong et al., 2003; Fariss et al., 1998; Kamei and Hollyfield, 1999). We found a significant (∼18-fold) increase in unglycosylated TIMP3 in SFD RPE ECM and a significant although smaller increase (∼4-fold) in the unglycosylated form. CRISPR-corrected SFD RPE resulted in significantly lower unglycosylated TIMP3 compared to uncorrected SFD RPE, although it was still higher than that of the normal control RPE. There may be additional modifiers that affect *TIMP3* expression in RPE that are yet unidentified. Despite high intracellular and ECM TIMP3 protein levels in SFD RPE, no overall change in MMP-2 and -9 inhibition was noted on zymography using cell culture media, and overall MMP inhibitory activity by TIMP3 was similar between SFD and control RPE. However, when corrected for TIMP3 protein expression, intracellular TIMP3 from SFD RPE was significantly less effective (∼5 fold) in inhibiting MMP activity compared to normal controls. Although there is a clear increase in intracellular and ECM TIMP3 by SFD RPE, recent findings indicate that SFD RPE culture media has less TIMP3 compared to controls as detected by ELISA (Hongisto et al., 2020). This suggests that evaluation of SFD mutant TIMP3 activity by zymography using culture media may not be reflective of overall TIMP3 activity, and investigation into SFD TIMP3 activity may be better served by assaying the activity of accumulated intracellular or ECM TIMP3.

Additional indirect evidence that there may be increased MMP activity in SFD RPE comes from targeted metabolomics showing that 4-hydroxyproline is the most significantly altered metabolite with a ∼2.7-fold increase (p-value ≤ 1.0 x 10 ) in SFD RPE compared to controls. 4-hydroxyproline is a collagen degradation product (Gordon and Hahn, 2010; Li and Wu, 2018; Shoulders and Raines, 2009), and its increased level in SFD RPE without a concomitant increase in ^13^C proline consumption suggests that elevated 4-hydroxyproline results from increased collagen degradation. Further, we found that SFD RPE cells have a 3-fold decrease in reduced glutathione, a critical metabolite in preventing damage from oxidative stress. That human SFD RPE are more vulnerable to oxidative stress complements recent findings that RPE in the SFD (S179C *TIMP3*) mouse model are in a pro-oxidant environment and are sensitive to low doses of an oxidizing agent (Wolk et al., 2020).

In conclusion, we found that SFD iPSC-RPE formed large sub-RPE deposits similar in composition to the basal laminar drusen found in SFD family member globes by immunofluorescence staining and TEM imaging. S204C *TIMP3* correction by CRISPR-Cas9 gene editing in SFD iPSC RPE resulted in significantly reduced basal laminar and sub-RPE calcium deposits. Targeted metabolomics showed that intracellular 4-hydroxyproline, a major breakdown product of collagen, is the most significantly elevated metabolite in SFD RPE, suggesting increased ECM turnover. Finally, SFD RPE cells have reduced levels of intracellular reduced glutathione and were found to be more vulnerable to oxidative stress. Together, our findings suggest that critical elements of SFD pathology can be demonstrated in culture and future studies using these *in vitro* models may lead to further insights into disease mechanisms.

## Acknowledgements

The authors wish to thank Ed Parker for assistance with TEM imaging, C. Dirk Keene for histology interpretation, Kelie Gonzalez for cell culturing, and Jessica Rowlan for assistance with sequencing. We also would like to thank Julie Mathieu, Chris Cavanaugh, and Jennifer Hesson in the Tom & Sue Ellison Stem Cell Core for assistance with CRISPR gene editing.

## Data Availability Statement

Data is reported in figures, materials and methods, or supplemental material. Additional information is available upon request from authors.

---

**Supplementary Figure 1. iPSC lines used in the study have normal karyotype.**

**Supplementary Figure 2. Correction of 610A>T (S204C) mutation in TIMP3 gene using CRISPR-Cas9 in SFD patient-derived iPSCs.** (A) Schematic diagram of the genome editing strategy for SFD iPSCs. The sgRNA is marked in red, and the PAM sequence is in blue. The cutting site of the CRISPR-Cas9 enzyme is indicated by a black arrow. The patient-mutated base in the green square was replaced with the wild type base, producing a single precise point mutation thus resulting in the repair of the disease-causing mutation. A silent mutation was introduced in the PAM region to prevent further cutting by Cas9, as indicated. The long sequence below is a single stranded DNA (ssDNA) donor with the desired repair mutations and homologous arms (45□bp on each side). Sanger sequencing results of TIMP3 in the SFD RD9 iPSC line harboring the 610A>T (p.Ser204Cys) mutation before (B) and after correction (C). (D) Sequencing of two potential off-target regions in both CRISPR-corrected SFD clones show no genetic editing.

**Supplementary Figure 3. Immunostaining of drusen components in iPSC-RPE and donor globes.** (A-G) Staining for TIMP3, ApoE, vitronectin, and clusterin was observed in BrM of normal globes. As expected, TIMP2 was located in the RPE and not in BrM. iPSC-RPE did not produce sub-RPE deposits of substantial size to be labeled by immunostaining. (G,H) Collagen VI was diffusely distributed throughout the thickened BrM in SFD globes and intensely in iPSC-RPE deposits. (I) Summary of human globe and iPSC-RPE immunostaining.

## References

Anand-Apte, B., Chao, J.R., Singh, R., Stohr, H., 2019. Sorsby fundus dystrophy: Insights from the past and looking to the future. J Neurosci Res 97, 88–97.

Arpino, V., Brock, M., Gill, S.E., 2015. The role of TIMPs in regulation of extracellular matrix proteolysis. Matrix Biol 44–46, 247-254.

Bailey, T.A., Alexander, R.A., Dubovy, S.R., Luthert, P.J., Chong, N.H., 2001. Measurement of TIMP-3 expression and Bruch’s membrane thickness in human macula. Exp Eye Res 73, 851–858.

Bonnans, C., Chou, J., Werb, Z., 2014. Remodelling the extracellular matrix in development and disease. Nat Rev Mol Cell Biol 15, 786–801.

Brydon, E.M., Bronstein, R., Buskin, A., Lako, M., Pierce, E.A., Fernandez-Godino, R., 2019. AAV-Mediated Gene Augmentation Therapy Restores Critical Functions in Mutant PRPF31(+/-) iPSC-Derived RPE Cells. Mol Ther-Meth Clin D 15, 392–402.

Buchholz, D.E., Pennington, B.O., Croze, R.H., Hinman, C.R., Coffey, P.J., Clegg, D.O., 2013. Rapid and efficient directed differentiation of human pluripotent stem cells into retinal pigmented epithelium. Stem Cells Transl Med 2, 384–393.

Carrero-Valenzuela, R.D., Klein, M.L., Weleber, R.G., Murphey, W.H., Litt, M., 1996. Sorsby fundus dystrophy. A family with the Ser181Cys mutation of the tissue inhibitor of metalloproteinases 3. Arch Ophthalmol 114, 737–738.

Chong, N.H., Alexander, R.A., Gin, T., Bird, A.C., Luthert, P.J., 2000. TIMP-3, collagen, and elastin immunohistochemistry and histopathology of Sorsby’s fundus dystrophy. Invest Ophthalmol Vis Sci 41, 898–902.

Chong, N.H., Kvanta, A., Seregard, S., Bird, A.C., Luthert, P.J., Steen, B., 2003. TIMP-3 mRNA is not overexpressed in Sorsby fundus dystrophy. Am J Ophthalmol 136, 954–955.

Curcio, C.A., 2012. Pathophysiology of Non-Neovascular Age-Related Macular Degeneration: The Oil Spill in Bruch’s Membrane and beyond. Ophthalmologica 228, 23–25.

Davis, W.L., Jones, R.G., Hagler, H.K., 1981. An electron microscopic histochemical and analytical X-ray microprobe study of calcification in Bruch’s membrane from human eyes. J Histochem Cytochem 29, 601–608.

Du, J., Yanagida, A., Knight, K., Engel, A.L., Vo, A.H., Jankowski, C., Sadilek, M., Tran, V.T., Manson, M.A., Ramakrishnan, A., Hurley, J.B., Chao, J.R., 2016. Reductive carboxylation is a major metabolic pathway in the retinal pigment epithelium. Proc Natl Acad Sci U S A 113, 14710–14715.

Fariss, R.N., Apte, S.S., Luthert, P.J., Bird, A.C., Milam, A.H., 1998. Accumulation of tissue inhibitor of metalloproteinases-3 in human eyes with Sorsby’s fundus dystrophy or retinitis pigmentosa. Br J Ophthalmol 82, 1329–1334.

Galloway, C.A., Dalvi, S., Hung, S.S.C., MacDonald, L.A., Latchney, L.R., Wong, R.C.B., Guymer, R.H., Mackey, D.A., Williams, D.S., Chung, M.M., Gamm, D.M., Pebay, A., Hewitt, A.W., Singh, R., 2017. Drusen in patient-derived hiPSC-RPE models of macular dystrophies. Proc Natl Acad Sci U S A 114, E8214–E8223.

Gliem, M., Muller, P.L., Mangold, E., Bolz, H.J., Stohr, H., Weber, B.H., Holz, F.G., Charbel Issa, P., 2015. Reticular Pseudodrusen in Sorsby Fundus Dystrophy. Ophthalmology 122, 1555–1562.

Gong, J., Cai, H., Team, N.G.S.C.A., Noggle, S., Paull, D., Rizzolo, L.J., Del Priore, L.V., Fields, M.A., 2020. Stem cell-derived retinal pigment epithelium from patients with age-related macular degeneration exhibit reduced metabolism and matrix interactions. Stem Cells Transl Med 9, 364–376.

Gordon, M.K., Hahn, R.A., 2010. Collagens. Cell Tissue Res 339, 247–257.

Hazim, R.A., Karumbayaram, S., Jiang, M., Dimashkie, A., Lopes, V.S., Li, D., Burgess, B.L., Vijayaraj, P., Alva-Ornelas, J.A., Zack, J.A., Kohn, D.B., Gomperts, B.N., Pyle, A.D., Lowry, W.E., Williams, D.S., 2017. Differentiation of RPE cells from integration-free iPS cells and their cell biological characterization. Stem Cell Res Ther 8.

Hogan, M.J., Alvarado, J.A., Weddell, J.E., 1971. Histology of the human eye : an atlas and textbook. Saunders, Philadelphia.

Hongisto, H., Dewing, J.M., Christensen, D.R.G., Scott, J., Cree, A.J., Nattinen, J., Maatta, J., Jylha, A., Aapola, U., Uusitalo, H., Kaarniranta, K., Ratnayaka, J.A., Skottman, H., Lotery, A.J., 2020. In vitro stem cell modelling demonstrates a proof-of-concept for excess functional mutant TIMP3 as the cause of Sorsby Fundus Dystrophy. J Pathol.

Jackson, H.W., Defamie, V., Waterhouse, P., Khokha, R., 2017. TIMPs: versatile extracellular regulators in cancer. Nat Rev Cancer 17, 38–53.

Jacobson, S.G., Cideciyan, A.V., Regunath, G., Rodriguez, F.J., Vandenburgh, K., Sheffield, V.C., Stone, E.M., 1995. Night blindness in Sorsby’s fundus dystrophy reversed by vitamin A. Nat Genet 11, 27–32.

Jin, M., Barron, E., He, S., Ryan, S.J., Hinton, D.R., 2002. Regulation of RPE intercellular junction integrity and function by hepatocyte growth factor. Invest Ophthalmol Vis Sci 43, 2782–2790.

Johnson, L.V., Forest, D.L., Banna, C.D., Radeke, C.M., Maloney, M.A., Hu, J., Spencer, C.N., Walker, A.M., Tsie, M.S., Bok, D., Radeke, M.J., Anderson, D.H., 2011. Cell culture model that mimics drusen formation and triggers complement activation associated with age-related macular degeneration. Proc Natl Acad Sci U S A 108, 18277–18282.

Kamei, M., Hollyfield, J.G., 1999. TIMP-3 in Bruch’s membrane: changes during aging and in age-related macular degeneration. Invest Ophthalmol Vis Sci 40, 2367–2375.

Khan, K.N., Mahroo, O.A., Khan, R.S., Mohamed, M.D., McKibbin, M., Bird, A., Michaelides, M., Tufail, A., Moore, A.T., 2016. Differentiating drusen: Drusen and drusen-like appearances associated with ageing, age-related macular degeneration, inherited eye disease and other pathological processes. Prog Retin Eye Res 53, 70–106.

Knupp, C., Chong, N.H., Munro, P.M., Luthert, P.J., Squire, J.M., 2002. Analysis of the collagen VI assemblies associated with Sorsby’s fundus dystrophy. J Struct Biol 137, 31–40.

Kuntz, C.A., Jacobson, S.G., Cideciyan, A.V., Li, Z.Y., Stone, E.M., Possin, D., Milam, A.H., 1996. Sub-retinal pigment epithelial deposits in a dominant late-onset retinal degeneration. Invest Ophth Vis Sci 37, 1772–1782.

Kuroda, T., Ando, S., Takeno, Y., Kishino, A., Kimura, T., 2019. Robust induction of retinal pigment epithelium cells from human induced pluripotent stem cells by inhibiting FGF/MAPK signaling. Stem Cell Res 39, 101514.

Li, P., Wu, G., 2018. Roles of dietary glycine, proline, and hydroxyproline in collagen synthesis and animal growth. Amino Acids 50, 29–38.

Pilgrim, M.G., Lengyel, I., Lanzirotti, A., Newville, M., Fearn, S., Emri, E., Knowles, J.C., Messinger, J.D., Read, R.W., Guidry, C., Curcio, C.A., 2017. Subretinal Pigment Epithelial Deposition of Drusen Components Including Hydroxyapatite in a Primary Cell Culture Model. Invest Ophthalmol Vis Sci 58, 708–719.

Qi, J.H., Anand-Apte, B., 2015. Tissue inhibitor of metalloproteinase-3 (TIMP3) promotes endothelial apoptosis via a caspase-independent mechanism. Apoptosis 20, 523–534.

Qi, J.H., Dai, G., Luthert, P., Chaurasia, S., Hollyfield, J., Weber, B.H., Stohr, H., Anand-Apte, B., 2009. S156C mutation in tissue inhibitor of metalloproteinases-3 induces increased angiogenesis. J Biol Chem 284, 19927–19936.

Sarks, S., Cherepanoff, S., Killingsworth, M., Sarks, J., 2007. Relationship of Basal laminar deposit and membranous debris to the clinical presentation of early age-related macular degeneration. Invest Ophthalmol Vis Sci 48, 968–977.

Shoulders, M.D., Raines, R.T., 2009. Collagen structure and stability. Annu Rev Biochem 78, 929–958.

Sonoda, S., Spee, C., Barron, E., Ryan, S.J., Kannan, R., Hinton, D.R., 2009. A protocol for the culture and differentiation of highly polarized human retinal pigment epithelial cells. Nat Protoc 4, 662–673.

Stohr, H., Anand-Apte, B., 2012. A review and update on the molecular basis of pathogenesis of Sorsby fundus dystrophy. Adv Exp Med Biol 723, 261–267.

Suzuki, M., Curcio, C.A., Mullins, R.F., Spaide, R.F., 2015. REFRACTILE DRUSEN: Clinical Imaging and Candidate Histology. Retina 35, 859–865.

Tan, A.C.S., Pilgrim, M.G., Fearn, S., Bertazzo, S., Tsolaki, E., Morrell, A.P., Li, M., Messinger, J.D., Dolz-Marco, R., Lei, J., Nittala, M.G., Sadda, S.R., Lengyel, I., Freund, K.B., Curcio, C.A., 2018. Calcified nodules in retinal drusen are associated with disease progression in age-related macular degeneration. Sci Transl Med 10.

Thompson, R.B., Reffatto, V., Bundy, J.G., Kortvely, E., Flinn, J.M., Lanzirotti, A., Jones, E.A., McPhail, D.S., Fearn, S., Boldt, K., Ueffing, M., Ratu, S.G., Pauleikhoff, L., Bird, A.C., Lengyel, I., 2015. Identification of hydroxyapatite spherules provides new insight into subretinal pigment epithelial deposit formation in the aging eye. Proc Natl Acad Sci U S A 112, 1565–1570.

Wang, L., Clark, M.E., Crossman, D.K., Kojima, K., Messinger, J.D., Mobley, J.A., Curcio, C.A., 2010. Abundant lipid and protein components of drusen. PLoS One 5, e10329.

Weber, B.H., Lin, B., White, K., Kohler, K., Soboleva, G., Herterich, S., Seeliger, M.W., Jaissle, G.B., Grimm, C., Reme, C., Wenzel, A., Asan, E., Schrewe, H., 2002. A mouse model for Sorsby fundus dystrophy. Invest Ophth Vis Sci 43, 2732–2740.

Weber, B.H., Vogt, G., Pruett, R.C., Stohr, H., Felbor, U., 1994. Mutations in the tissue inhibitor of metalloproteinases-3 (TIMP3) in patients with Sorsby’s fundus dystrophy. Nat Genet 8, 352–356.

Wolk, A., Upadhyay, M., Ali, M., Suh, J., Stoehr, H., Bonilha, V.L., Anand-Apte, B., 2020. The retinal pigment epithelium in Sorsby Fundus Dystrophy shows increased sensitivity to oxidative stress-induced degeneration. Redox Biol 37, 101681.

Zhang, Q., Chrenek, M.A., Bhatia, S., Rashid, A., Ferdous, S., Donaldson, K.J., Skelton, H., Wu, W., See, T.R.O., Jiang, Y., Dalal, N., Nickerson, J.M., Grossniklaus, H.E., 2019. Comparison of histologic findings in age-related macular degeneration with RPE flatmount images. Molecular vision 25, 70–78.

