## Supplemental Figures and Tables for "Extracellular Matrix Dysfunction in Sorsby Patient-Derived Retinal Pigment Epithelium"

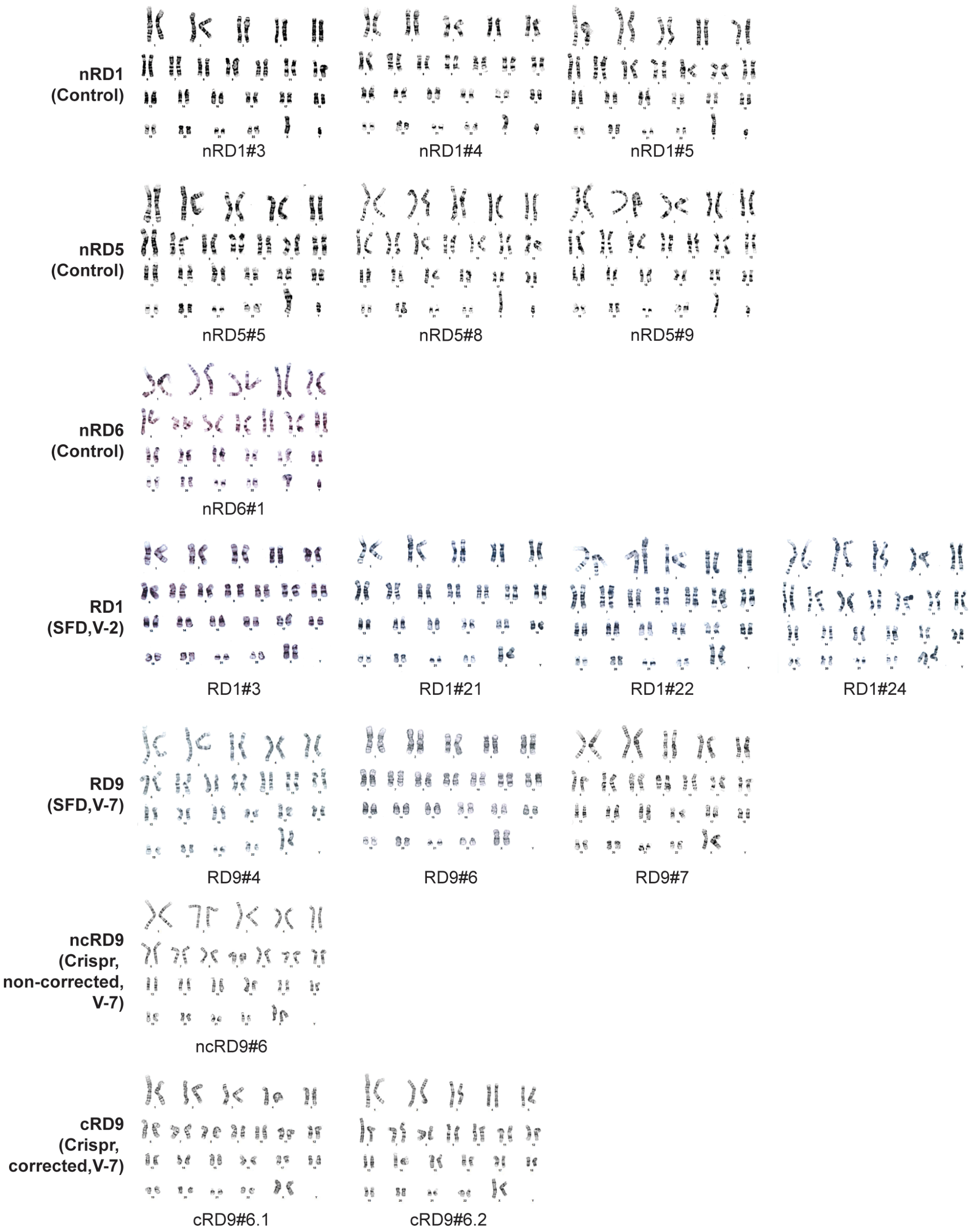


### Supplementary Figure 1

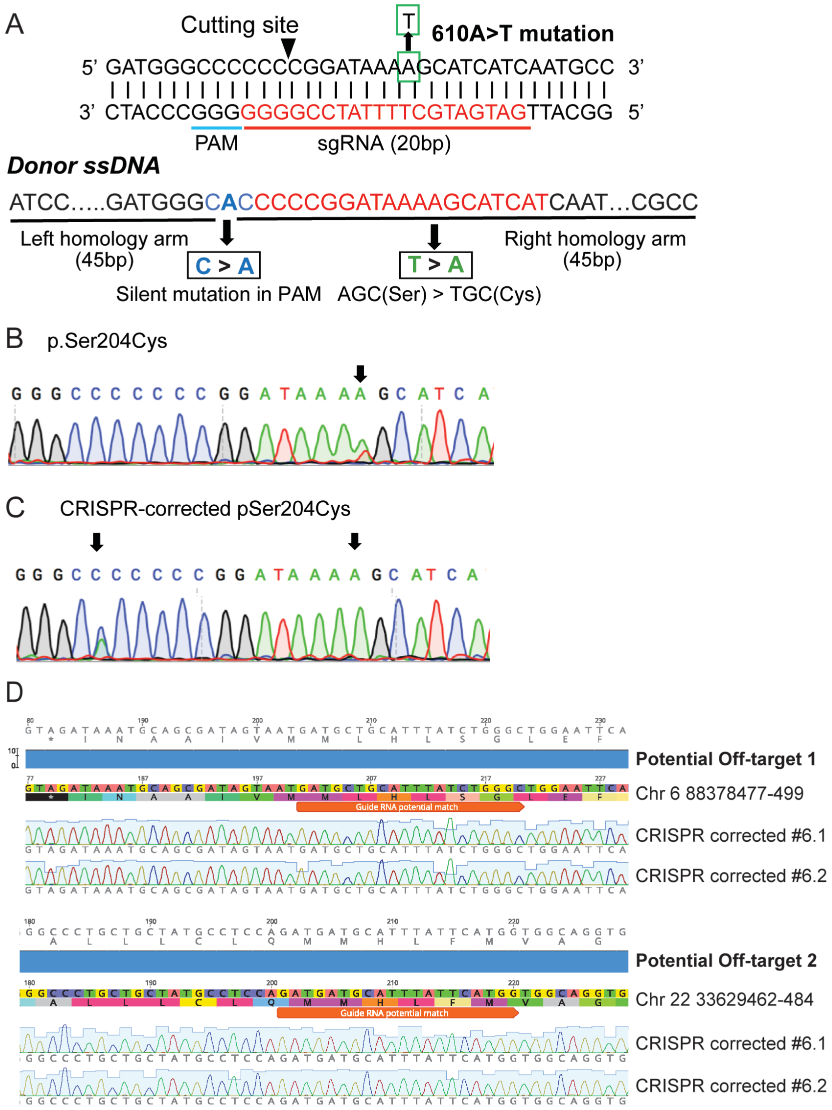


### Supplementary Figure 2

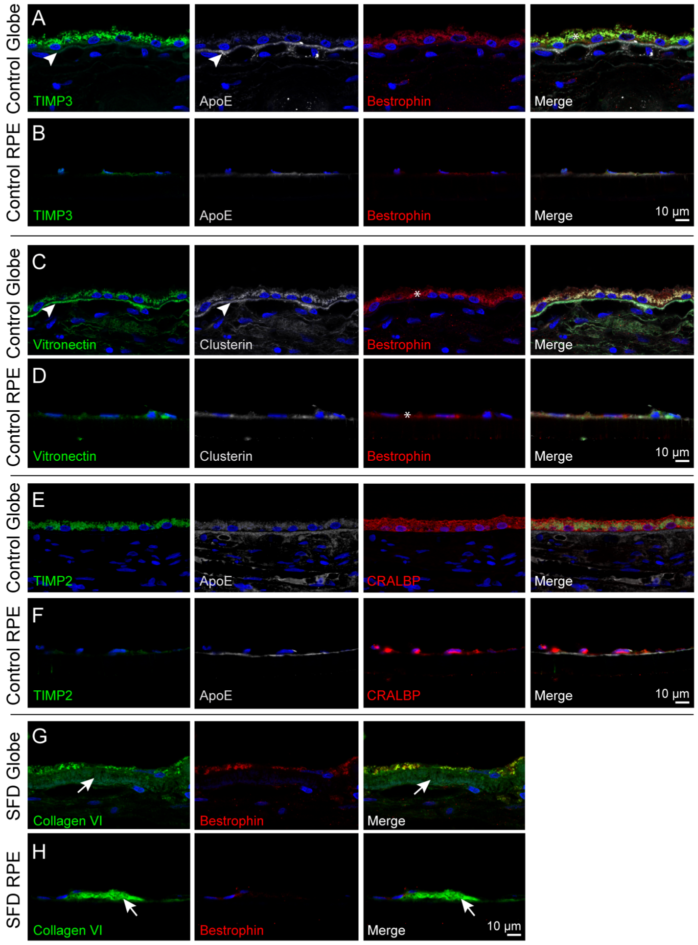


| Marker | Control Globe | S204C Globes | Control iPSC RPE | SFD iPSC RPE |
|---|---|---|---|---|
| APOE | RPE: +<br>BrM: ++ | RPE: +<br>sub-RPE deposit: ++ | RPE: + | RPE: ++<br>sub-RPE deposit: ++ |
| Clusterin | RPE: ++<br>BrM: ++ | RPE: +<br>sub-RPE deposit: ++ | RPE: ++ | RPE: +<br>sub-RPE deposit: +++ |
| Collagen VI | RPE: ++<br>BrM: ++ | RPE: +++ (speckled)<br>sub-RPE deposit: ++ | RPE: ++ | RPE: ++<br>sub-RPE deposit: +++ |
| TIMP2 | RPE: +++<br>BrM: None | RPE: ++<br>sub-RPE deposit: None | RPE: + | RPE: ++<br>sub-RPE deposit: None |
| TIMP3 | RPE: +++<br>BrM: + | RPE: +++<br>sub-RPE deposit: ++ | RPE: + | RPE: ++<br>sub-RPE deposit: ++ |
| Vitronectin | RPE: ++<br>BrM: ++ | RPE: +<br>sub-RPE deposit: +++ | RPE: + | RPE: +<br>sub-RPE deposit: ++ |

**Supplementary Figure 3**

Supplementary Table 1

| | Identifier | Age/ Sex | ARMS2 | | CFH | EFEMP1 | HTRA1 | TIMP3 | | |
|---|---|---|---|---|---|---|---|---|---|---|
| | | | A69S (G>T) | indel (polyA tail) | Y402H (T>C) | R345W (C>T) | G>A | S204C (A>T) | T191C (A) | S179C (C) |
| **CONTROL** | nRD1 | 77/M | Hetero | indel | Homo | Normal | Hetero | Normal | Normal | Normal |
| | nRD5 | 69/M | Hetero | indel | Homo | Normal | Hetero | Normal | Normal | Normal |
| | nRD6 | 66/M | Normal | Normal | Normal | Normal | Normal | Normal | Normal | Normal |
| **SFD** | RD1 | 66/F | Homo | indel | Hetero | Normal | Homo | Hetero | Normal | Normal |
| | RD9 | 58/F | Hetero | indel | Homo | Normal | Hetero | Hetero | Normal | Normal |

Supplementary Table 2

| RPE Cell Lines | iPSC Clone/ RPE # | Figures |
|---|---|---|
| nRD1 (Control) | #2 | 2D- F, 6G |
| | #3 | 5D |
| | #4 | 2B, 3C&D, 6C |
| | #5 | 2B&C;3A,3C&D,5A,5D |
| | #6 | 2B, 6F |
| nRD5 (Control) | #5 | 2B- F, 5B-D, 6B&C, 6G, Sup Fig 3B&D, 3F, 3I |
| | #8 | 2D, 3C&D, 5C, 6G |
| | #9 | 2D-F, 5A, 5C, 6G |
| nRD6 (Control) | #1 | 2A, 2C-F, 5B, 6G |
| RD1 (SFD, V-2) | #21 | 2B-F, 3C&D, 5A,5C&D,6C, E&G |
| | #22 | 2B-F, 3C&D, 5B-D, 6E&G |
| | #24 | 2B-F, 3C&D, 5A-D, 6E&G |
| RD9 (SFD, V-7) | #4 | 2B-F, 3C&D, 5B-D, 6E&G, Sup Fig 3I |
| | #6 | 2B-F, 3B, 3C&D, 3E-E", 4B, 4D, 4F, Sup3H&I, 5A, 5C-D, 6B&C, 6E&G |
| | #7 | 2A, 2B-F, 3C&D, 5A&B, 5D,6E&G, Sup Fig 3I |
| ncRD9 (CRISPR, uncorrected, V-7) | #1 | 6F&G |
| cRD9 (CRISPR, corrected, V-7) | #6.1 | 6B,C,F,G |
| | #6.2 | 6B,C,D,E,G |

Supplementary Table 3

**Primers sequences used in sequencing**

| *Gene* | *Mutation* | *Forward (5'-3')* | *Reverse (5' – 3')* |
|---|---|---|---|
| ARMS2 | A69S | *GTTCCTGTGTCCTTCATTTCCACTC* | *GGTAAGGCCTGATCATCTGCATT* |
| ARMS2 | Insertion-deletion | *GCCACACTGTGCAACCTGGAATTTC* | *AGGTTGTCCTCTAGATCCAGGGTG* |
| CFH | Y402H | *GTGCAAACCTTTGTTAGTAACTTTAGTTCG* | *CAAGGTGACATAAACATTTTGCCAC* |
| EFEMP1 | R345W | *GTGTTGGCATAAATAGTTGTGGCCTGTA* | *AGGCAGAGTCAGTGGTTAACATTGGG* |
| HTRA1 | Promoter G>A | *CATCTGAGACCGCTCCACCCTCGCCAGTTA* | *CGCTTTCCGCAGGTTCTCTCGCTGAGATTC* |
| TIMP3 | S179C / T191C / S204C | *GTAGCCTCAGGCCTGGGCATA* | *TGTCATCATCTGGGAAGAGTTAGTGTC* |
| TIMP3 | Off-Target 1 | *AGTAGACCCTGACAATGCCACG* | *TCAAAACCCTAACATAACTTTGGCC* |
| TIMP3 | Off-Target 2 | *CCATTCTTTCAAGGATCGTTTATCTTTCTC* | *AACTTTGTCTTGGCTTCAGGGTTCAGAGA* |

**Primers sequences used in qPCR**

| *Gene* | *Forward (5'-3')* | *Reverse (5' – 3')* |
|---|---|---|
| ApoE | *CTGGCACTGGGTCGCTTTT* | *AGTTGTTCCTCCAGTTCCGATTT* |
| Bestrophin | AACCATTGGAAACATTTAACTCAGAC | AGTGCCGTTGTTCAGTTCTAATC |
| COL6A1 | AGCAAGTGTGCTGCTCCTTC | CTTCCAGGATCTCCGGCTTC |
| CRALBP | CACGCTGCCCAAGTATGATG | CCAGGACAGTTGAGGAGAGG |
| GAPDH | TGAAGGTCGGAGTCAACGGA | CCATTGATGACAAGCTTCCCG |
| RPE65 | TACAGAAAGCACTGAGTTGAGC | CCATTTAGTAAGTCCACATTCATTTCC |
| TIMP3 | GCAGATAGACTCAAGGTGTGTGAAA | TCCCTCACTCTTACATGCAGACA |
| Vitronectin | GAAGCCGTCAGAGATATTTCG | CCTTCACCGACCTCAAGAAC |

Supplementary Table 4

| METABOLITE | CAS | PLATFORM | POLARITY | PRECURSOR (Da) | PRODUCT (Da) |
|---|---|---|---|---|---|
| 3-hydroxybutyric acid | 300-85-6 | LC MS/MS | Positive | 105 | 23 |
| 3-hydroxykynurenine | 484-78-6 | LC MS/MS | Negative | 223 | 75 |
| a-ketoglutarate | 328-50-7 | LC MS/MS | Negative | 145 | 101 |
| Alanine | 02-72-7 | LC MS/MS | Negative | 134 | 107 |
| AMP | 61-19-8 | LC MS/MS | Negative | 346 | 134 |
| Ascorbic Acid | 50-81-7 | LC MS/MS | Negative | 175 | 87 |
| Asparagine | 70-47-3 | LC MS/MS | Positive | 133 | 70 |
| Aspartate | 56-84-8 | LC MS/MS | Positive | 134 | 74 |
| ATP | 56-65-5 | LC MS/MS | Positive | 508 | 136 |
| Carnitine | 541-15-1 | LC MS/MS | Positive | 163 | 85 |
| cGMP | 7665-99-8 | LC MS/MS | Negative | 344 | 150 |
| Choline | 62-49-7 | LC MS/MS | Positive | 104 | 60 |
| Citrate | 126-44-3 | LC MS/MS | Negative | 191 | 87 |
| Creatinine | 60-27-5 | LC MS/MS | Negative | 112 | 41 |
| FAD | 146-14-5 | LC MS/MS | Positive | 786 | 136 |
| Fumarate | 142-42-7 | LC MS/MS | Positive | 117 | 59 |
| Glutamate | 11070-68-1 | LC MS/MS | Positive | 148 | 84 |
| Glutamine | 56-85-9 | LC MS/MS | Positive | 147 | 84 |
| Glycine | 56-40-6 | LC MS/MS | Positive | 76 | 30 |
| GSH (glutathione) | 70-18-8 | LC MS/MS | Negative | 306 | 143 |
| GSSG (oxidized glutathione) | 121-24-4 | LC MS/MS | Negative | 611 | 306 |
| GTP | 86-01-1 | LC MS/MS | Negative | 522 | 159 |
| Hydroxyproline | 51-35-4 | LC MS/MS | Positive | 132 | 86 |
| Isoleucine | 443-79-8 | LC MS/MS | Positive | 132 | 69 |
| Lactate | 50-21-5 | LC MS/MS | Negative | 89 | 43 |
| Leucine | 61-90-5 | LC MS/MS | Positive | 132 | 86 |
| Lysine | 56-87-1 | LC MS/MS | Positive | 147 | 84 |
| Methionine | 63-68-3 | LC MS/MS | Positive | 150 | 61 |
| N1-Methylnicotinamide | 3106-60-3 | LC MS/MS | Positive | 137 | 78 |
| NAD | 53-84-9 | LC MS/MS | Positive | 664 | 136 |
| NADH | 58-68-4 | LC MS/MS | Negative | 664 | 397 |
| NADP | 53-59-8 | LC MS/MS | Positive | 744 | 136 |
| Nicotinamide | 98-92-0 | LC MS/MS | Positive | 123 | 80 |
| Nicotinic acid | 59-67-6 | LC MS/MS | Negative | 122 | 78 |
| Ophthalmic acid | 495-27-2 | LC MS/MS | Positive | 290 | 58 |
| Phenylalanine | 63-91-2 | LC MS/MS | Positive | 166 | 120 |
| Proline | 147-85-3 | LC MS/MS | Positive | 116 | 70 |
| Pyroglutamic acid | 98-79-3 | LC MS/MS | Positive | 130 | 84 |
| Pyruvate | 57-60-3 | LC MS/MS | Negative | 87 | 43 |
| Serine | 56-45-1 | LC MS/MS | Positive | 106 | 60 |
| Succinate | 56-14-4 | LC MS/MS | Negative | 117 | 73 |
| Taurine | 107-35-7 | LC MS/MS | Positive | 126 | 108 |
| Threonine | 72-19-5 | LC MS/MS | Positive | 120 | 102 |
| Trigonelline | 535-83-1 | LC MS/MS | Positive | 138 | 92 |
| Tyrosine | 60-18-4 | LC MS/MS | Positive | 182 | 136 |
| Valine | 72-18-4 | LC MS/MS | Positive | 118 | 72 |
